# AI for Fisheries Science: Neural Network Tools for Forecasting, Spatial Standardization, and Policy Optimization

**DOI:** 10.64898/2026.03.13.711664

**Authors:** Maia S. Kapur, Grant Adams, Marcus Lapeyrolerie, James T. Thorson

**Affiliations:** NOAA Fisheries, Alaska Fisheries Science Center, Seattle, WA 98105; University of British Columbia, Department of Forest Resources Management, Vancouver, BC V6T 1Z4

**Author notes:** Corresponding author: Maia Kapur. **Data Availability:** Code to obtain and/or re-simulate data for and reproduce the case studies is available at https://github.com/mkapur/deep-fish.

## Abstract

The development of Artificial Intelligence (AI) presents novel opportunities for tackling complex marine resource management challenges. Among AI models, neural networks are a powerful class of tools capable of learning nonlocal and lagged patterns from fisheries data as well as approximating nonlinear relationships among multiple latent variables using estimation methods that automatically implement statistical shrinkage. This gives them potential to effectively handle data obtained from fisheries populations subject to dynamic environments. We highlight two flexible subclasses and one application of neural networks: Long Short-Term Memory (LSTM) and Convolutional Neural networks (CNNs) and policy discovery via Reinforcement Learning. LSTMs are designed to handle sequential data by allowing prediction from past values at both short and long time-lags. CNNs are not explicitly designed to handle temporal information, but can interpolate a spatial latent variable based on its value within a geographic neighborhood, and can be combined with LSTM models for this purpose. This “Food for Thought” paper introduces and applies these neural network approaches, both alone and in combination, to demonstrate their potential application for several foundational topics in fisheries science: 1) the forecasting of population weight-at-age, 2) the standardization of spatio-temporal indices of relative abundance, and 3) the discovery of harvest policies to optimize catches and maintain spawning biomass. Each section provides a simple, simulated example and describes the tradeoffs – particularly the lack of inferential capability – presented by using neural networks over traditional approaches for each topic. We then outline medium-term research questions that may clarify, facilitate or qualify the applicability of these tools to fisheries management science. Finally, we discuss how future combinations of these approaches could result in simplified ways to estimate and forecast stock biomass and provide harvest advice.

## Introduction

### Motivation

Science based fisheries management is currently facing unprecedented challenges due to shifting funding priorities and dynamic and unexpected changes in the marine environment (Patrick and Link, 2015). These challenges are co-occurring with an expansion of methodologies designed to address the diverse data types now available for some fisheries, from molecular information informing movement and demographic estimation (Punt et al., 2024) to fine-scale acoustic monitoring data (Griffin et al., 2018). However, fisheries data have traditionally been analyzed using biological models fit statistically to data, which are constrained by, potentially biased, model assumptions. Artificial intelligence (AI) tools are less constraining in their assumptions and have been explored for data collection and preliminary analysis (such as the statistical post-stratification of fisheries data (Gasper and Cahalan, 2025, p. 202), occurrence records (Morand et al., 2024) or automated image detection (Saqib et al., 2024), but the broader potential of AI remains underexplored. There are simultaneous efforts to modernize the modelling infrastructure used to assess fishery populations (Maunder et al., 2025), presenting an opportune moment to revisit the methodological landscape of fisheries stock assessment. This paper focuses on neural network models as a promising avenue to modernize scientific fisheries management in the coming decades. We provide readers the core concepts of neural network models, in contrast to familiar concepts from generalized linear modeling and machine learning and share three demonstrative case studies.

### Pointwise Regression: A Common Tool in Fisheries Management

Most existing fisheries management modeling approaches rely on regression and parametric process based models. These include traditional approaches such as generalized linear models (GLMs) through many machine learning (ML) methods, which include boosted regression trees, random forests, projection pursuit regression, lasso, and Gaussian process models (to name a few; Hastie et al., 2001), and has been used in fisheries science for over a decade (Rubbens et al., 2023). ML and GLMs are similar in that they both typically define a pointwise regression, where each sample is predicted by a vector of associated covariates; these can include complex methods of handling spatio-temporal processes, such as tinyVAST (Thorson et al., 2025). Due to this conceptual similarity between ML and GLMs, there have been many previous analyses in fisheries science comparing ML methods with conventional GLMs or hierarchical models (Stock et al., 2020), or extending regression models to include a Gaussian process component (Sugeno and Munch, 2013; Thorson et al., 2014). These methods have the advantage of being easy to implement and facilitate inference, but are limited by assumptions regarding the statistical distribution of modeled data and are sensitive to mis-specification.

### Neural Networks: an overview of a foundational AI technique

Neural networks are a class of machine learning models inspired by the structure and function of the mammalian brain. They are constructed using a series of transformations of features and penalized model weights that can approximate any function. This architecture offers advantages for handling high-dimensional, nonlinear and spatiotemporal data types common in fisheries; their value to ecology for this reason has been recognized for nearly three decades (Lek et al., 1996). In some cases, neural networks can outperform traditional process-based or statistical models in predicting ecological processes. Alternatively, neural networks can be embedded within process based models to improve performance (e.g., (Triebe et al., 2021; Wesselkamp et al., 2024)).

We claim in the following that neural networks (NNs) are often more suited than previous ML methods to identify complex features that are present in fisheries analysis because of their ability to learn nonlinear and complex patterns. Neural networks penalize complexity differently from GLMs by the progressive updating of model weights (Fan et al., 2021), and can optionally set a subset of model weights to zero to prevent overfitting (Srivastava et al., 2014). These approaches are known as “implicit” and “explicit” regularization, respectively. Usefully, these methods for regularization do not require marginalizing across any coefficients, and are therefore much faster than regularization in Bayesian or empirical-Bayes hierarchical models. Current NN methods have also seen less use in fisheries science; one aim of the present paper is to promote the exploration of these methods. Table 1 provides an explicit comparison between familiar concepts in statistical-ecology and equivalent (though not always identical) concepts in neural networks/artificial intelligence

**Table 1.** Non-exhaustive correspondence of vocabulary from statistical ecology to neural network/AI methods.

| Term in statistical ecology | Term in neural networks | Brief description |
| --- | --- | --- |
| deterministic latent variable | layer | A hidden but non-random state predicted from the model |
| nonlinear transformation for element of a latent variable | neuron | A method of abstraction to map unobserved effects to responses, for example, through logistic or exponential curves |
| reverse-mode automatic differentiation | backpropagation | Algorithm to efficiently compute gradients by working backwards through the model |
| parameter | weight | A model value controlling the strength of an input's effect |
| parameter estimation | training | The process of adjusting parameters or weights to minimize error on observed data |
| regression modelling | supervised learning | Predicting outcomes from inputs using observed data |
| gradient-based optimizer<sup>2</sup> | Stochastic gradient descent (e.g., Adam) | Methodology for updating model parameters during training to minimize error |
| optimizing a management policy | stochastic policy ascent | A process of adjusting policy parameters to optimize a specific reward function |
| numerical overflow and underflow | Exploding or vanishing gradient | Because the gradient calculation involves products of terms, and each term in the product might be very large or small, the gradient might |

<sup>2</sup> Where each optimization step is based on a subset of data; hyperparameters control how the explored parameters update given the gradient, and each loop through all partitions of the data is called an “epoch”

|  |  |  |
| --- | --- | --- |
|  |  | end up being smaller or larger than the numerical minimum or maximum allowed. This can preclude accurate calculation of the gradient. |

### Potential applications of Neural Networks in Fisheries

There are numerous types of neural network models, and this paper focuses on two for their potential suitability to the spatial and/or temporal dynamism common to marine fisheries. Recurrent Neural Networks (RNNs) are a subtype of neural network designed to handle sequential data such as time series using feedback loops that maintain an internal state or memory of previous inputs in the series. This recurrent structure enables RNNs to capture temporal dependencies and model the evolution of dynamical systems. They are natural candidates for nonlinear autoregressive models, where future values are predicted based on past values.

However, basic RNNs can suffer from the numerical underflow or overflow issues (see Table 1) during training, which makes it difficult for them to learn long-range dependencies in the data. Long Short-Term Memory (LSTM) networks are a specialized type of RNN architecture designed to mitigate these issues. LSTMs introduce a cell state, which acts as a long-term memory, and sub-states known as “gates” that control the flow of information into and out of the cell state, allowing the network to selectively remember and forget information over long sequences. The value of RNNs like LSTMs lies in their ability to model and predict the behavior of partially observed dynamical systems, where not all relevant state variables are directly measured. By learning the temporal patterns in the observed data, RNNs can make predictions about future states, which is highly valuable in fisheries science for forecasting population dynamics, catch, and other time-dependent variables.

Convolutional Neural Networks (CNNs) were developed for the computer vision field as a technique to facilitate the detection and labeling of images. In the simplest case, this involves a multi-step process of passing (“convolving”) a set of learnable convolutional filters, or kernels, to iteratively extract patterns. This process builds a hierarchical numerical representation, often referred to as a feature map or embedding, that captures high-level features at low spatial resolution. This allows the CNN to effectively learn spatial dependencies and patterns in the image, which could be a photograph of an animal or a map of observed biomass from a fishery independent survey, presenting an intriguing possibility for the standardization of spatially-explicit data. However, the basic CNN framework is not designed to explicitly handle temporal data nor irregular or sparse datasets characteristic of most fisheries surveys or catch time series. Researchers have combined CNN and LSTM neural network approaches to produce forecasts of spatial processes (e.g., Yang et al., (2025); the authors are aware of a single example wherein CNN and LSTM neural network approaches were combined to produce forecasts of probable catches (Agmata and Guðmundsson, 2025), although other studies have used CNN in isolation (Morand et al., 2024). Crucially, that example did not conduct an explicit comparison between the proposed CNN+LSTM approach, variations thereof, and currently-used methods for handling spatio-temporal data, such as design-based expansion or regression-based standardization tools such as tinyVAST, which we do in this case study.

### Aims and Structure of this Paper

This “Food for Thought” paper introduces and applies these neural network approaches, both alone and in combination, as a forward-looking demonstration for several foundational topics in fisheries science: 1) the forecasting of population processes, with size-at-age as an example; 2) the standardization of spatio-temporal indices of relative abundance, and 3) the discovery of harvest policies to optimize catches and maintain spawning biomass. For each case study we describe the methods used to produce the illustrative example, and discuss the relevance for the fisheries assessment and management audience. Where possible, we provide direct mapping between existing concepts in statistical ecology (such as spatio-temporal standardization or interpolation) or fisheries management (such as management strategy evaluation, MSE; for this reason, the section on reinforcement learning for management is longer than the others). Advancing these techniques could enhance the adaptability, precision and sustainability of scientific fisheries management in a rapidly changing environment. The paper concludes with a “Research Roadmap” outlining the most important research questions to be considered in evaluating the tractability of these tools for fisheries management in the coming decades.

#### Case Studies

This paper presents three case studies. Each is designed to highlight how NNs may be applied to a variety of data types and processes central to fisheries management science. We also selected these to represent a spectrum of tractability and impact, ranging from applications that could be implemented today to those that would require completely novel management frameworks Figure 1. We emphasize that the case study methods are examples and are not final; more work is needed by the fisheries science community to investigate the tradeoffs, specifications, and nuances of each application before they are ready for operational use.

**Figure 1.**
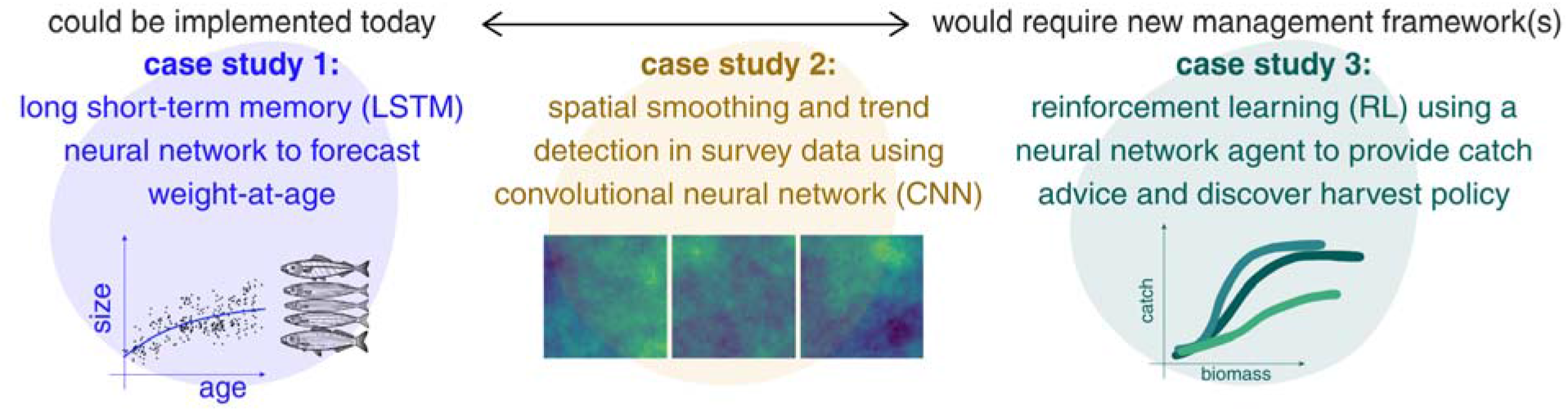
Conceptual diagram of case studies. The case studies were selected and are ordered (from left to right) in terms of how tractably each NN application could be included in current scientific fishery management frameworks.

#### Case Study 1: Neural Networks for Stock Demography Forecasting

Even highly complex stock assessment models typically resort to simplification of observed demographic processes in order to make management decisions. In particular, projections of future size-at-age form the basis for fisheries management advice, yet scientists will commonly use a historical or recent (e.g. 5-year) mean of observed fish sizes, though size-at-age can change dramatically among years (Stawitz et al., 2019). Stock assessment models also rely on estimates of historical size-at-age to derive estimates of historical biomass. Process-based models, such as the von-Bertalanffy growth curve, are commonly used for estimating average size-at-age. Given that size-at-age is well-observed for many managed fisheries, is sensitive to stochastic environmental processes, and underpins catch advice for industrial fisheries management, this case study illustrates how NN can provide flexibility to predict observable processes in fisheries.

We specifically compared the performance of two NN approaches to several existing methods for deriving size-at-age. The models included a simple average of size-at-age from the terminal five years of data (‘mean-5’), a three parameter von-Bertalanffy curve (‘VBGF’); a three parameter von-Bertalanffy curve with IID year univariate random effects on asymptotic size and growth rate (VBGF-RE’); a von Bertalanffy curve with 3D AR1 year, age, and cohort random effects (‘GMRF’; Cheng et al., 2023); a simple 3-layer NN (‘NN’), and a LSTM (‘LSTM’); see Supplementary Material S.1. for further details..

To explore whether the relative performance of the different approaches depend on the temporal structure of the simulation, we fit all six models to two simulated size-at-age datasets with parameters based on Bering Sea pollock (*Gadus chalcogrammus*) from NOAA’s Alaska Fisheries Science Center. The first simulated dataset simulated data from a three parameter von-Bertalanffy curve representing constant time-invariant growth. The second simulated dataset simulated data from a three parameter von-Bertalanffy curve with time-varying parameters following an AR1 and directional trend representing time-varying growth. Model parameters and sample sizes used for simulated data-sets were based on sampling effort and life history parameters of cod like species. We produced 300 30-year replicates of each dataset. Only simulations in which all models converged were retained.

We evaluated prediction performance on each dataset by calculating the average root mean squared error (RMSE) across ages from a 10-fold cross validation where random years of data were removed from each fold. One- and two-year projection performance was evaluated using five retrospective forecast peels. This allowed us to compare average RMSE across peels, both for hindcast and projection accuracy and between the time-invariant or time-dependent scenarios.

This simulation exercise demonstrated that the LSTM most frequently resulted in the lowest RMSE for all three prediction periods (hindcast, one and two years forward) for the time-invariant growth simulations (Figure 2). When simulated size-at-age varied through time, average RMSE was most frequently lowest for the GMRF model for the hindcast and one-year projection (Figure 2), though the LSTM and mean-5 performed comparably or better for the two-year projection in this scenario.

**Figure 2.**
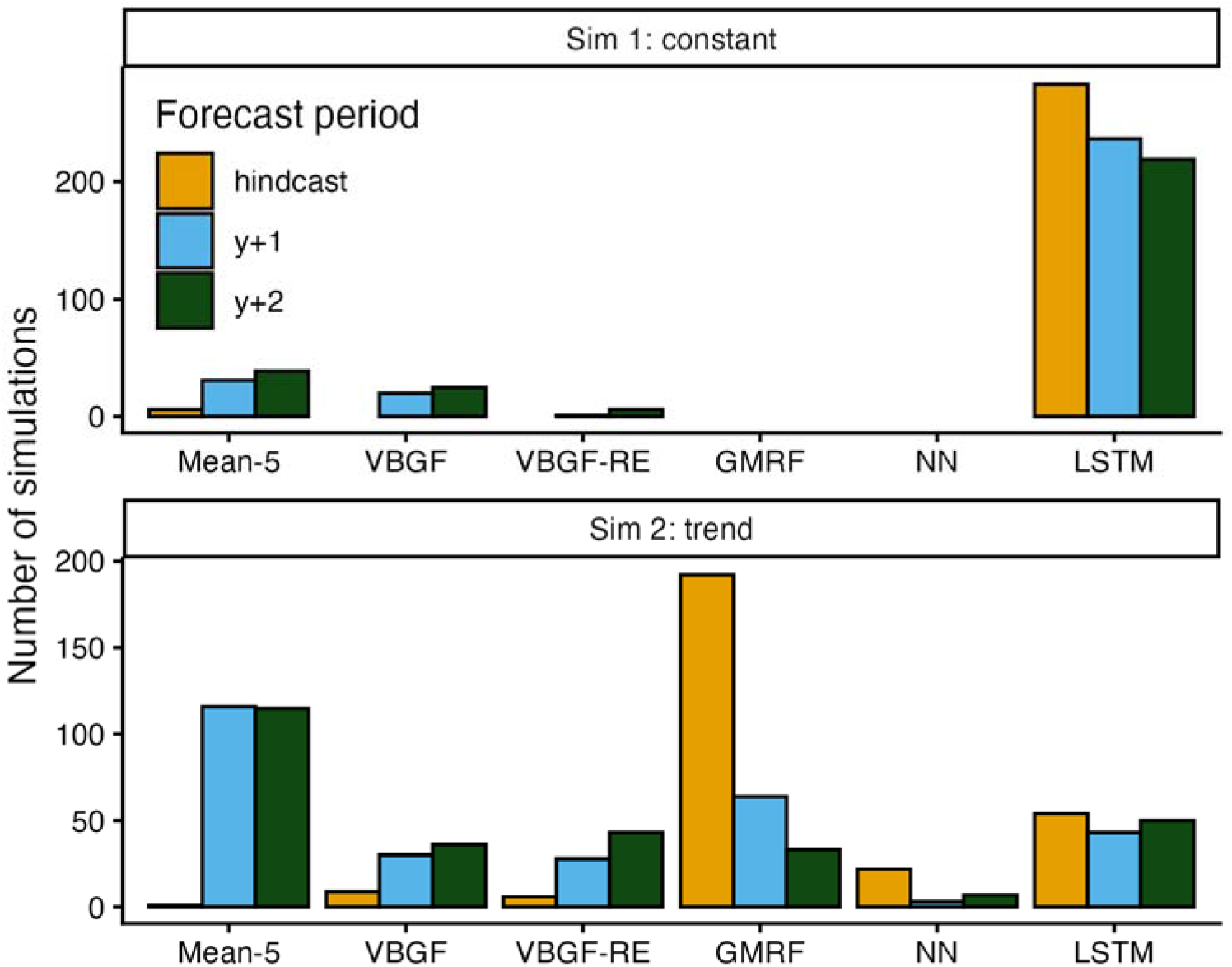
Case Study 1: Number of times each process- and neural network based model resulted in the lowest average root mean squared error (RMSE) when fit to simulated data (n = 300) without (top row) or with (bottom row) a temporal trend in true fish growth. Hindcast represents 10-fold leave year out cross-validation and “y+1” and “y+2” represents forecast skill from sequential peels of historical data and forecasted for two future years.

The GMRF is designed to handle temporal variation explicitly and is able to separate observation uncertainty from the latent temporal variability; it is possible that with lower observation error the performance of the LSTM method would have been improved. For both size-at-age scenarios, we also visualized how predictive performance varied across ages, which had a less distinct pattern (Supplementary Figures S1 and S2).

#### Case Study 2: Standardizing survey data through space and time using a convolutional neural network

Several methods have been developed to model or standardize for spatio-temporal processes in fisheries data, particularly survey observations. These methods arose from the recognition that underlying population processes and the methods for observing fish populations are subject to variation in space and time, and accounting for this variation is required to ensure unbiased estimates of population size or trajectory. The tinyVAST framework (Thorson et al., 2025) builds upon the Vector Autoregressive Spatio-Temporal (VAST, Thorson (2019)) modeling approach that allows users to specify separable Gaussian Markov Random fields and delta-GLMMs. The latest approach allows the specification of a broader class and flexibility of multivariate models including spatial factor analysis and ARIMA (Jenkins, 2014) structures, enabling the representation of simultaneous, lagged and recursive dependencies common to ecological and fishery processes. TinyVAST is similar to sdmTMB (Anderson et al., 2022) in that they both utilize the modern Template Model Builder language (Kristensen et al., 2016), incorporate SPDE-based spatial precision matrices, though the former is better suited for allowing for multivariate temporal dependencies. Surprisingly, the recognition that neural network models are useful for data with strong spatial dependencies (Wikle and Zammit-Mangion, 2023) has not yet led to rigorous investigation of how these techniques compare to existing approaches for standardizing biomass observations into annual indices of fishery abundance.

The simulation study was conducted on a 20 x 20 spatial grid over 12 years, with spatial correlation modeled through a row-standardized neighborhood matrix (rho = 0.95) and temporal correlation through an AR1 process (p = 0.8). The underlying simulated biomass included both spatial and temporal random effects; in each year 200 cells (50% of the domain) were randomly sampled for observation that arose from a Tweedie distribution (Zainol Mustafa and Nadia, 2025; rho = 1.5, psi = 0.2). The experiment was replicated 25 times to assess model performance and stability. For each replicate, observed data were passed to two CNN-based models (described below) as well as to tinyVAST to a) interpolate continuous spatio-temporal biomass estimates and b) calculate annual indices of abundance. A design-based estimator was included for comparison for the annual indices. Design-based expansion is a simple calculation that takes the average observation across sampled cells and multiplies it across the entire domain based on the fraction of observed cells; this precludes the production of continuous surfaces but facilitates comparison of annual indices.

We included two versions of the CNN approach, which differed in their handling of missing data and temporal dependencies. Both approaches relied on the network to learn spatial relationships through coordinate embeddings and produced continuous surfaces of estimated biomass. The simpler approach, labeled “CNN” in results, took spatial coordinates as input and processed each year separately; only points with observed (sampled) data were included in training. These vectorized inputs passed through two dense embedding layers (dimensions 64 and 128, with swish activation), reshaped to a 16×4×2 spatial tensor and processed by two 3×3 two-dimensional convolutional layers with ReLU activation. The second approach, labeled “CNN-LSTM” explicitly handled sparsity in the observed data via a binary masking channel which informed the model where observations existed in a given year. Temporal information in the CNN-LSTM was incorporated via a lag-based approach, whereby predictions at each timestep were informed by the previous three years of data. Both CNN models were implemented in R using the keras3 package (Kalinowski et al., 2025), compiled with Adam and a trainable Tweedie loss function with sigmoid power parameter initialized to 1.5 and constrained to (1, 2) specified to support comparison with tinyVAST. The 12-year, 25-repliate experiment to fit the three models took 16 hours to run on a personal computer. Performance was compared across models by examining the trajectory of standardized indices, and overall RMSE in log-biomass and log-density (across all replicates and years).

This simulation exercise demonstrated that neither of the CNN-based models out-performed the existing tinyVAST approach in terms of log-biomass and log-density, though all approaches were able to produced standardized indices of similar scale to the true biomass (Figure 3). recent work has replicated our finding that CNNs can match, but not surpass, traditional species distribution modeling approaches (Kellenberger et al., 2026). Examination of the interpolated map for a single year and replicate shows that while both CNN-based models were able to identify areas of relatively higher and lower biomass only tinyVAST was able to recover small-scale regional patterns. This application of CNNs certainly extends beyond the traditional implementation, wherein an intact image (or collection of images for training) would be passed to the model. CNNs are known to perform poorly when images are incompletely observed (Heinke et al., 2021), and are also known to perform poorly when training data are limited (most CNNs are trained on thousands of images, not 12 years of data as in this case)

**Figure 3.**
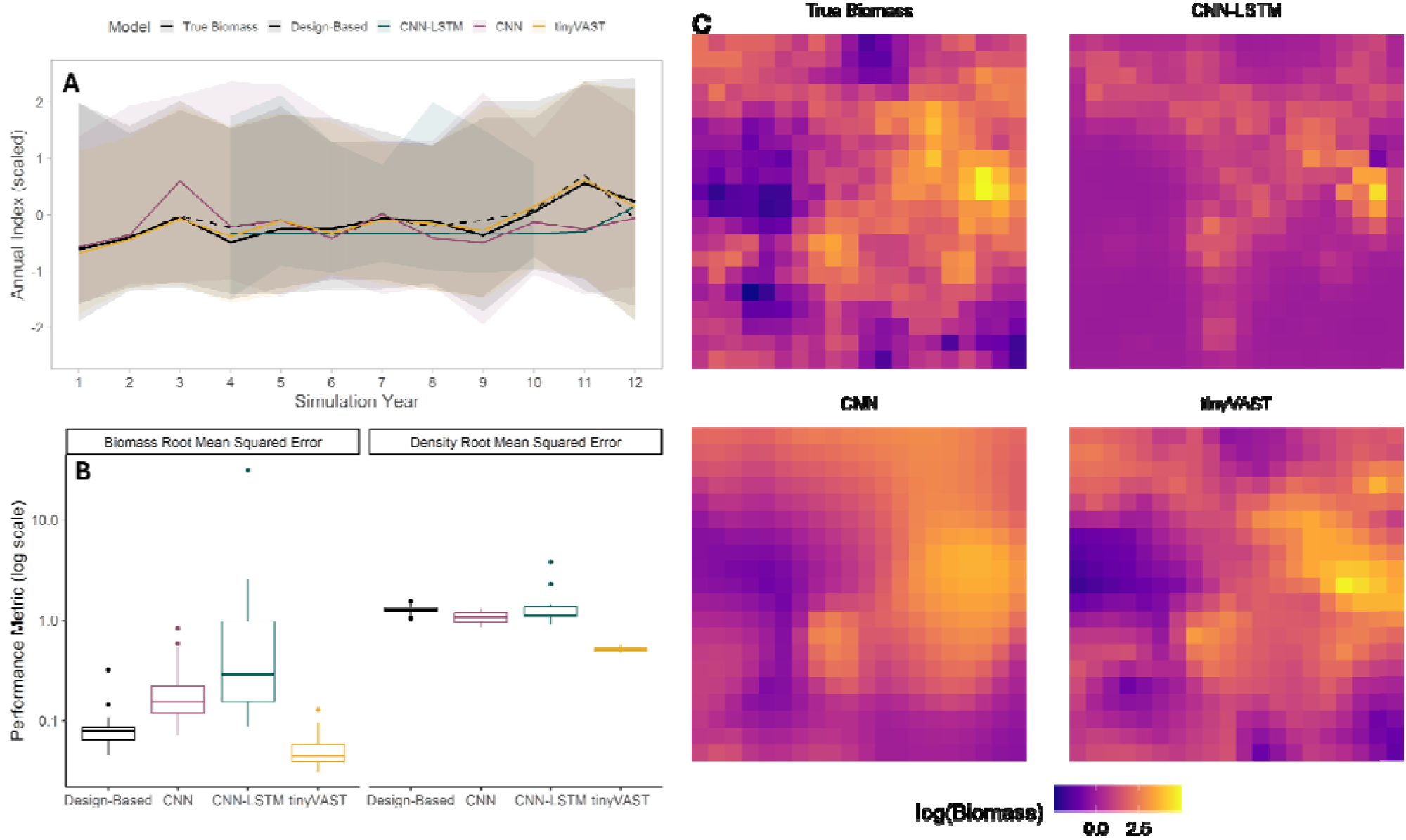
Case Study 2: Investigation of neural networks for spatio-temporal survey data interpolation and index standardization. A) Mean scaled annual indices (lines) and 95% confidence interval (shaded ribbon) for 25 replicates of a simulated 12-year biomass time series; black solid line is true biomass whereas all other colors are model estimates. B) Performance metrics of various estimation models (colors) in terms of RMSE in biomass (lefthand figure) or density (righthand side) for estimates in a 20×20 gridded domain for 12 simulated years across 25 replicates. C) Maps of true and estimated log biomass across the domain for a single replicate and year; results shown only for models that produce an interpolated surface as part of the estimation step.

**Figure 4.**
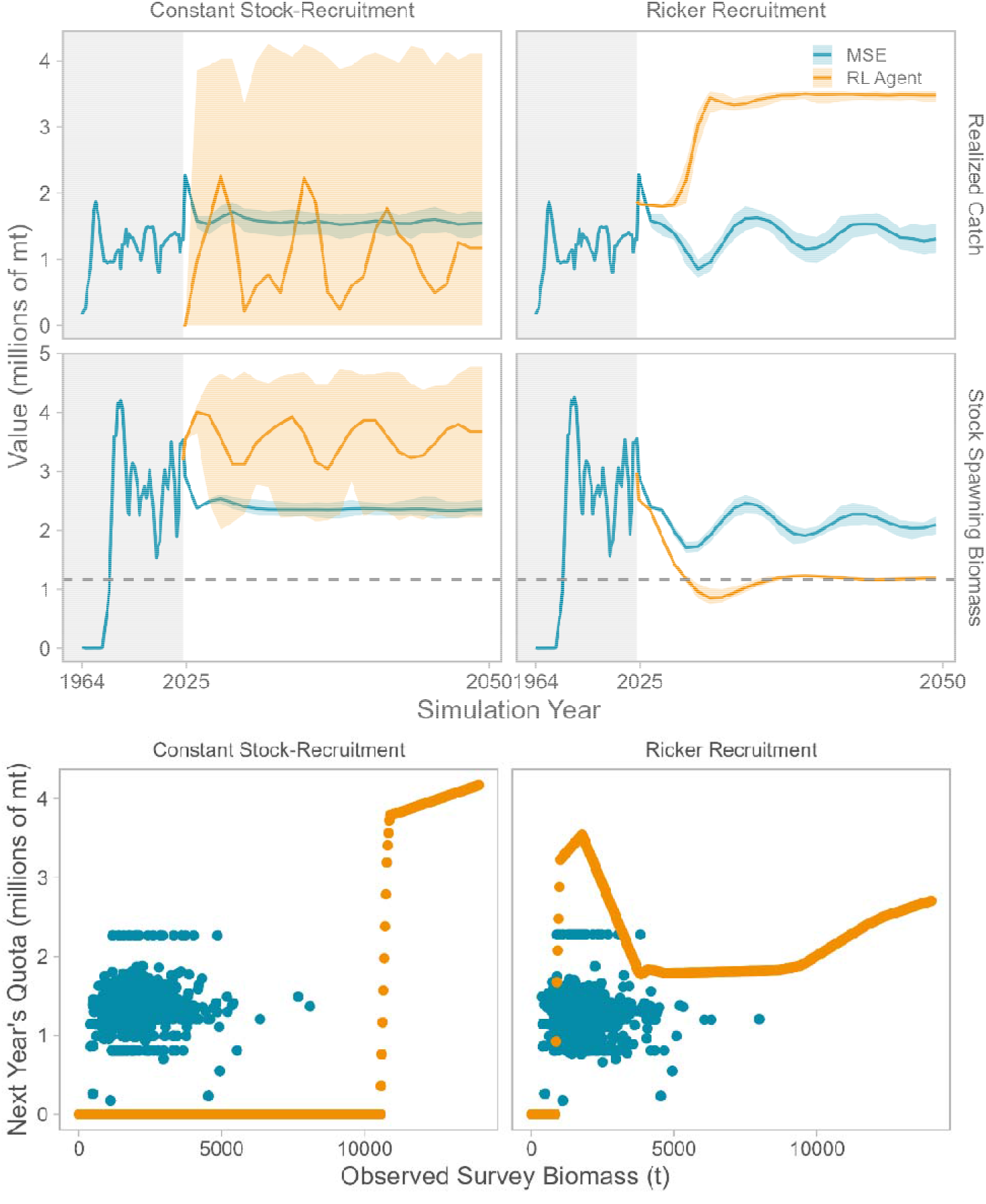
Case Study 3: Investigation of neural-network based reinforcement learning method (orange lines and points) for fisheries management, in comparison to 25 replicates of an MSE (blue lines and points) for EBS pollock using an OM conditioned on observations from 1964-2024 (grey rectangle; years are compressed) with a projection period from 2025-2050. Top row: realized catch (millions of mt) under the constant (1a) and Ricker (1b) recruitment scenarios. Middle row: stock spawning biomass in the operating model under the constant (2a) and Ricker (2b) recruitment scenarios. Horizontal dashed line indicates 20% of unfished biomass and the colored line represents the median across simulation replicates for a given year. Colored ribbons represent 95% simulation interval. Bottom row: realized harvest policy in terms of the quota (millions of mt) specified for future years(s) (points) versus the observed Bottom Trawl Survey Biomass (t) arising from each method under the constant (3a) or Ricker (3b) recruitment scenarios. The MSE did not use a survey-based method for setting quotas, but observations from 25 MSE replicates and the subsequent years’ catch are shown for comparison.

There are promising applications of neural networks that could be used for the detection of spatial patterns, some of which are in active development (e.g., (Deng et al., 2025)) and would likely require the generation of continuous spatio-temporal loadings from observed data. However, this case study indicates that tinyVAST is an appropriately precise and efficient method for the straightforward task of index standardization using sparse data; more sophisticated extensions to the CNN approach such as a vision transformer (Dosovitskiy et al., 2021) could be revisited for well-sampled fishery populations, and/or from interpolated meshes derived from such data.

### Case Study 3: Setting next year’s catch using reinforcement learning (RL)

A common problem is fisheries is the selection of management strategies to achieve policy objectives. Management strategy evaluation is a simulation framework that was developed to evaluate alternative strategies under uncertainty. Traditional MSE is a closed-loop forward-in-time simulation framework used to evaluate the performance of harvest strategies under uncertainty that can include an operating model (OM), estimation model (EM) and management rule. MSE is inherently sequential and forward-looking, as it mimics real-world decision-making over time. Crucially, the mechanism (or ‘policy’) by which future catches are determined is nearly always a predefined rule which is not informed or modified by the simulation itself. This means it is not tractable to explore all possible policies using traditional MSE. MSE is a time-consuming process that requires stakeholder input (Goethel et al., 2019) and the manual specification of alternative states of nature, estimation models, and harvest policies (Punt et al., 2016). Recent work has highlighted that static reference points like B_0_ (unfished biomass, the foundation of many management systems) are highly sensitive to model assumptions (Edgar et al., 2024) and fundamentally problematic for data-poor or dynamic fisheries to the degree that empirical data streams may offer more robust guidance for management in such situations (Blamey et al., 2025). These issues justify the exploration of data-driven methods, such as reinforcement learning, that can discover policies autonomously.

Reinforcement learning is a subfield of machine learning that is dedicated to solving sequential decision-making problems and consists of two primary components: the agent and the environment. The RL agent is a trainable neural network that interacts repeatedly with the ‘environment’, a representation of the population dynamics akin to the operating model used in Management Strategy Evaluation (Butterworth, 2007; Punt et al., 2016). During this interactive process (called “training”) the agent receives a feedback signal known as the “reward^1^” which it seeks to optimize by updating a policy, analogous to a management rule in MSE. A key distinction between MSE and RL is that the policy in RL is updated according to the agent’s experience in the environment, whereas the HCR in MSE remains static across simulations.

This case study directly compared the performance of a traditional management strategy evaluation with a RL-derived policy for the Eastern Bering Sea (EBS) pollock fishery. This allowed us to investigate the tractability and relative performance of the RL approach for a fishery with complex dynamics (multiple fishing fleets, age-structure) and a high degree of data availability (including weight-at-age and compositional data), yet using a simple, singular objective (maximization of annual harvest). This case study therefore extends beyond recent contributions that focus on simplified or simulated stock dynamics (e.g., Montealegre-Mora et al. (2025)). Two recruitment scenarios were explored for both approaches: the first utilized a stock independent approach where expected recruitment was defined by random annual deviations around a constant value (‘Constant recruitment’), and the second utilized a Ricker stock-recruitment relationship, where stock output varied both following random annual deviations and as a function of stock size (‘Ricker recruitment’). For EBS pollock, recruitment dynamics are a noted uncertainty for this stock and both parameterizations have been explored for management advice (Ianelli et al., 2024). This allowed us to determine whether commonly-used recruitment dynamics impacted relative performance or stability of the RL approach for this fishery. We compared future performance of the catch recommendations determined by the MSE and RL approaches by examining the median simulation trajectory and uncertainty of stock spawning biomass realized catches during the projection period. For illustration, we also compared the harvest policy learned by the RL approach to the survey observations and realized catches from the MSE. Full specifications of this case study can be found in Supplementary Material section 2.

The reinforcement learning (RL) agent trained under constant recruitment developed a harvest policy resembling precautionary harvest control rules. It avoided fishing when bottom trawl survey biomass was below about 10,500 mt and increased harvest sharply to ~4 million mt above that threshold, reaching ~4.25 million mt at higher levels. This produced large year-to-year fluctuations—ranging from zero to nearly double historical catches—but consistently maintained spawning biomass above that of the management strategy evaluation (MSE) scenario, which had stable catches (~1.5 million mt) and biomass (~2.5 million mt).Under the Ricker recruitment scenario, the RL agent’s policy was non-linear but smoother than in the constant recruitment case. It maintained nearly constant catches (1.8–3 million mt) for survey observations above 4,000 t, unexpectedly increased catches around 2,500 t, and stopped fishing below 1,000 t. This strategy stabilized spawning biomass near 1.25 million mt (~21% of unfished levels; 20%B_0_ is a common cutoff for fishery closure) while sustaining harvests of ~3.5 million mt—substantially higher than the MSE scenario, which averaged 1–1.5 million mt of catch and 2–2.5 million mt of spawning biomass. Despite its unconventional shapes, the RL policy outperformed the MSE in cumulative yield while maintaining more stable trajectories under the more complex Ricker dynamics.

These results expand upon findings by Montealegre-Mora (2025) to suggest that the RL approach is capable of discovering harvest policies for complex, data-rich fisheries and produce catch and spawning biomass trajectories of similar magnitude to those obtained by MSE under two distinct recruitment scenarios. That study demonstrated that RL can provide insights into HCR design that conventionally used methods in fisheries management are unable to achieve. This is especially intriguing given that the RL did not need to “step through” an updated estimation model at each timestep, presenting a potential companion or alternative to the time-consuming process of updating stock assessments in resource-limited settings.

Future work could explore constraints on the policy curve (i.e., forcing recti-linear policies or setting strict upper limits on catches, though these can inhibit agent training). The RL method could also be extended to base the current state of the population on more data types (age composition data, for instance) or to include representations of observation or process error (e.g., via curriculum learning, see Table 2). Finally, it would be useful to explore comparison of the RL method to empirical harvest control rules that set catches based on recent survey observations, particularly in the context of data quality and availability. The RL method employed here assumes that annual survey observations are equally representative of the population among years and through time; it would be useful to know if there are lower limits to the frequency and precision of survey observations that facilitate RL learning.

**Table 2.** Comparison of key concepts in reinforcement learning-based policy discovery and Management Strategy Evaluation.

|  | **Reinforcement Learning for Fisheries** | **Management Strategy Evaluation** |
| --- | --- | --- |
| **Representation of population/system dynamics** | Operating model conditioned to historical data, termed “environment” | Operating model conditioned to historical data |
| **Data source used for model selection** | Future simulation or historical data | Always future simulation |
| **Representation of future stochasticity** | Simulated variation in future recruitment deviations | Simulated variation in future population processes, typically recruitment deviations |
| **Representation of observation and/or estimation uncertainty** | Can be addressed with specialized neural network architectures (i.e., environmental uncertainty as hidden state) | Handled explicitly by estimation model |
| **How catches are selected** | The policy function, which is learned from direct interactions with the environment, prescribes the annual catch according to the observed state | Pre-defined harvest control rule sets annual catches based on the current state (which may be one or more estimated quantities or observed data) |
| **How performance is measured** | reward function is maximized during training as the agent interacts with the environment; performance metrics can be calculated after training based on the operating model time series, | After simulations, calculate performance metrics using operating model time series during projection period |
|  | as in MSE |  |
| **Comparison across policies** | Harvest policies compared internally as part of learning. Novel harvest control rules explored and evaluated; can consider distinct yet flexible rules; no consideration of estimation models | Can consider distinct yet pre-defined harvest control rules and/or estimation models as strategy components |

#### Summary of findings from Case Studies

Our case studies highlight the flexibility that neural networks present in addressing foundational topics in fisheries science, their immediate shortcomings, and directions for future research. For the prediction of weight-at-age, it appears valuable to include LSTM-based approaches as part of an analytical pipeline to evaluate forecast and hindcast projection accuracy given the frequency with which they out-performed other methods in terms of RMSE. For some prediction categories, the GMRF method showed similar performance in the presence of temporal variation in growth. For the development of annual indices of abundance from spatio-temporal survey data, or interpolation of biomass in space, neither the CNN nor CNN-LSTM approaches appeared to outperform tinyVAST. It is important to recognize that the CNN technique was developed for computer-vision applications under the assumption that the entire image is available to the network; the sparse information provided by our simulated survey was insufficient for the CNN to accurately characterize spatial dependencies, is precisely what tinyVAST was designed to handle. The RL-to-MSE comparison exercise highlighted promising potential for novel policy discovery, with the tradeoff that absent strong guidance, policies might be un-intuitive (hindering stakeholder communication and support) or induce untenably large variation in population trajectories.

#### Food for Thought: Priority Research Questions for AI in Fisheries Science

When statistical catch-at-age software tools such as Stock Synthesis became widespread, the fisheries science community spent nearly two decades examining the impacts of ‘misspecification’ on model performance, highlighting how un-accounted-for dynamics in spatial structure, fish growth, selectivity patterns, and more could bias the estimation of management quantities and subsequent advice. Now the fisheries science community must apply that same focus to uncovering the trade-offs and biases presented by using neural networks in tandem with our existing process-based modeling workflows.

Here, we pose several outstanding questions regarding the potential use of neural network models for fisheries science, which we categorize based on their use in projections, for setting quotas, or for fisheries science and research (Table 3). In particular, fisheries process models were designed to represent the mechanisms that underly population responses to fishing (Beverton and Holt, 1957). By representing these mechanisms, fisheries scientists presumably hoped to develop a “structural causal model” (Pearl, 2009), which could then be used to predict fishery responses to policy decisions that have not previously been seen (i.e., the simultaneous impact of fishing and climate change). We therefore recommend specific comparison of existing process-based and new neural-network models regarding their ability to predict (and quantify uncertainty) about fishery responses to previously unobserved policies; fishery scientists may need to expand their vocabulary for describing sources of uncertainty to include those common in the AI literature (for example, Allen Akselrud, 2024). Our first two case studies emphasized “predictive performance,” and good predictive performance is no guarantee of good performance for disentangling multiple causes when recommending new policies (Arif and MacNeil, 2022). Given the large literature regarding causal modelling in artificial intelligence research (reviewed briefly in Luo et al., 2020), we are optimistic that future neural network research can develop robust fisheries policies using monitoring data for systems involving multiple drivers.

**Table 3:**
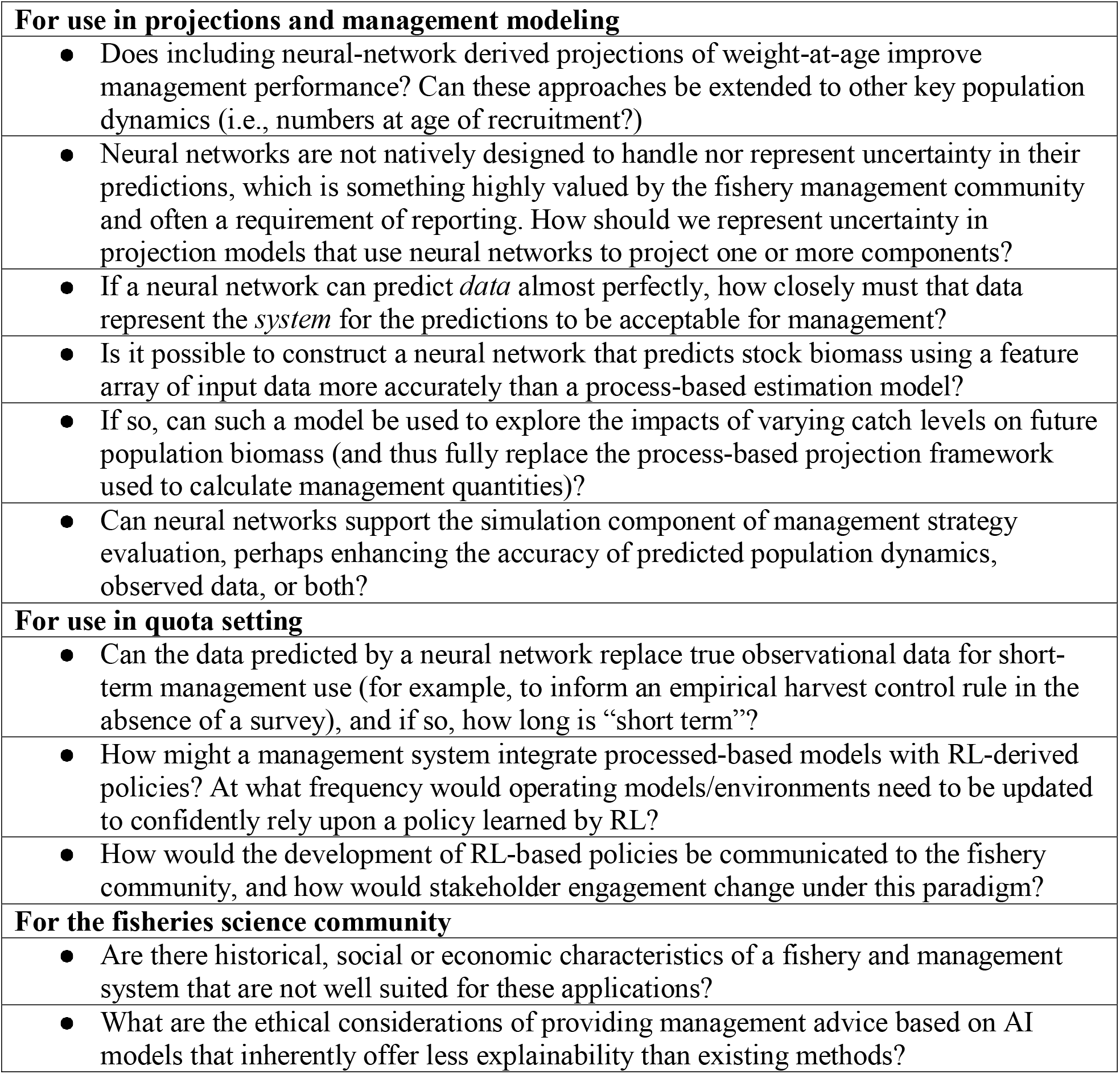
List of questions arising from case studies, that could form the basis of future research in scientific fisheries management.

## Conclusion

This Food-for-thought paper sought to 1) introduce the scientific fishery management community to modern techniques in deep learning with neural networks, exploring specific applications to foundational fisheries topics. Our case studies highlight that these methods have promise to supplement, improve upon, or change the ways we manage industrial fisheries, and underscore that rigorous benchmarking (comparison to existing methods) should be employed as they are further explored and refined.

## Footnotes

1 For readers seeking a broader introduction to RL in the context of ecology, see Lapeyrolerie et al., (2022).

