## Supplementary Material for "AI for Fisheries Science: Neural Network Tools for Forecasting, Spatial Standardization, and Policy Optimization"

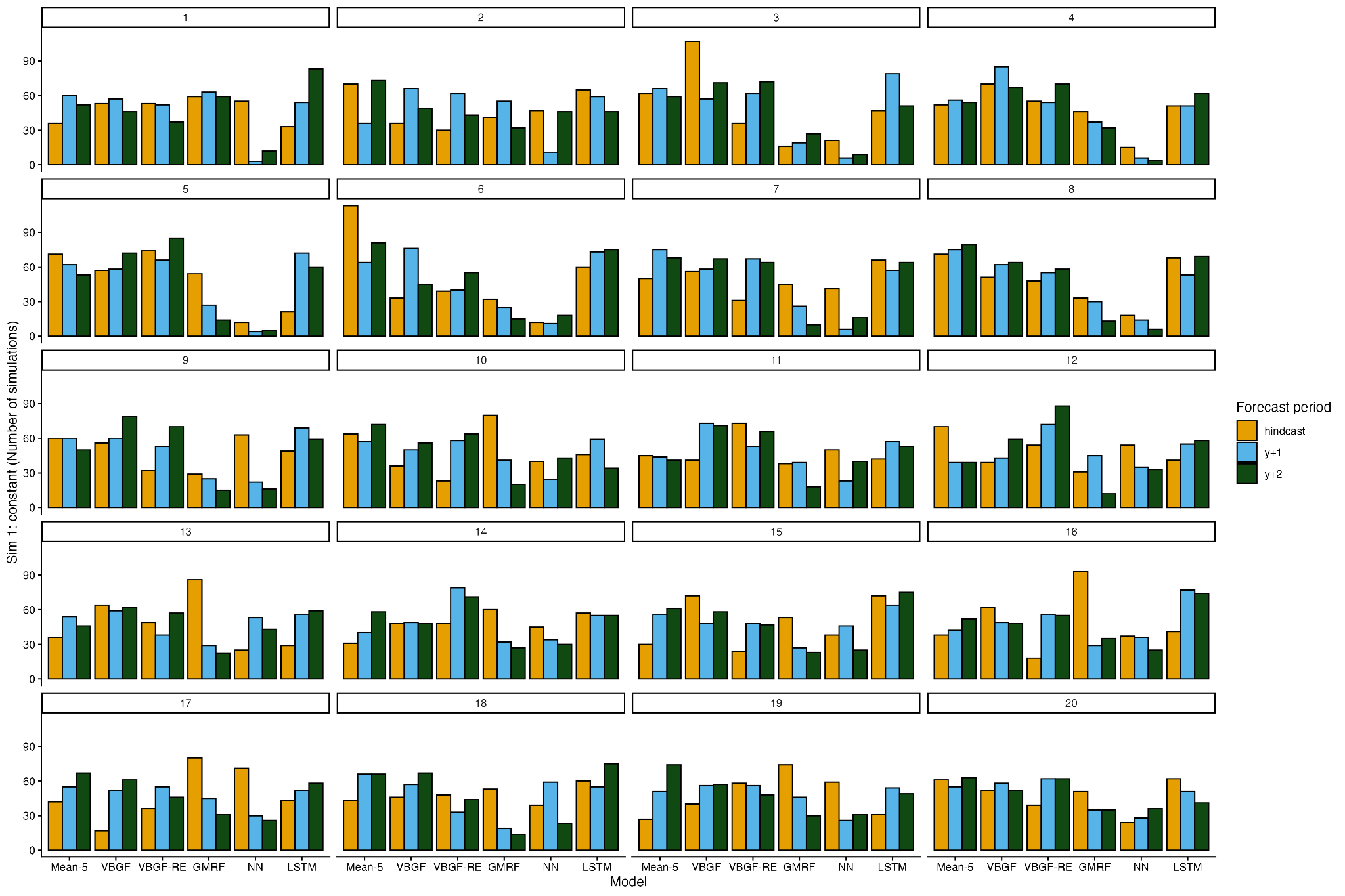

**Figure S1**. Case Study 1: Number of times each process- and neural net based model resulted in the lowest average root mean squared error (RMSE) per age when fit to time-invariant simulated data (n = 300). Hindcast represents 10-fold leave year out cross-validation and “y+1” and “y+2” represents forecast skill from sequential peels of historical data and forecasted for two future years.

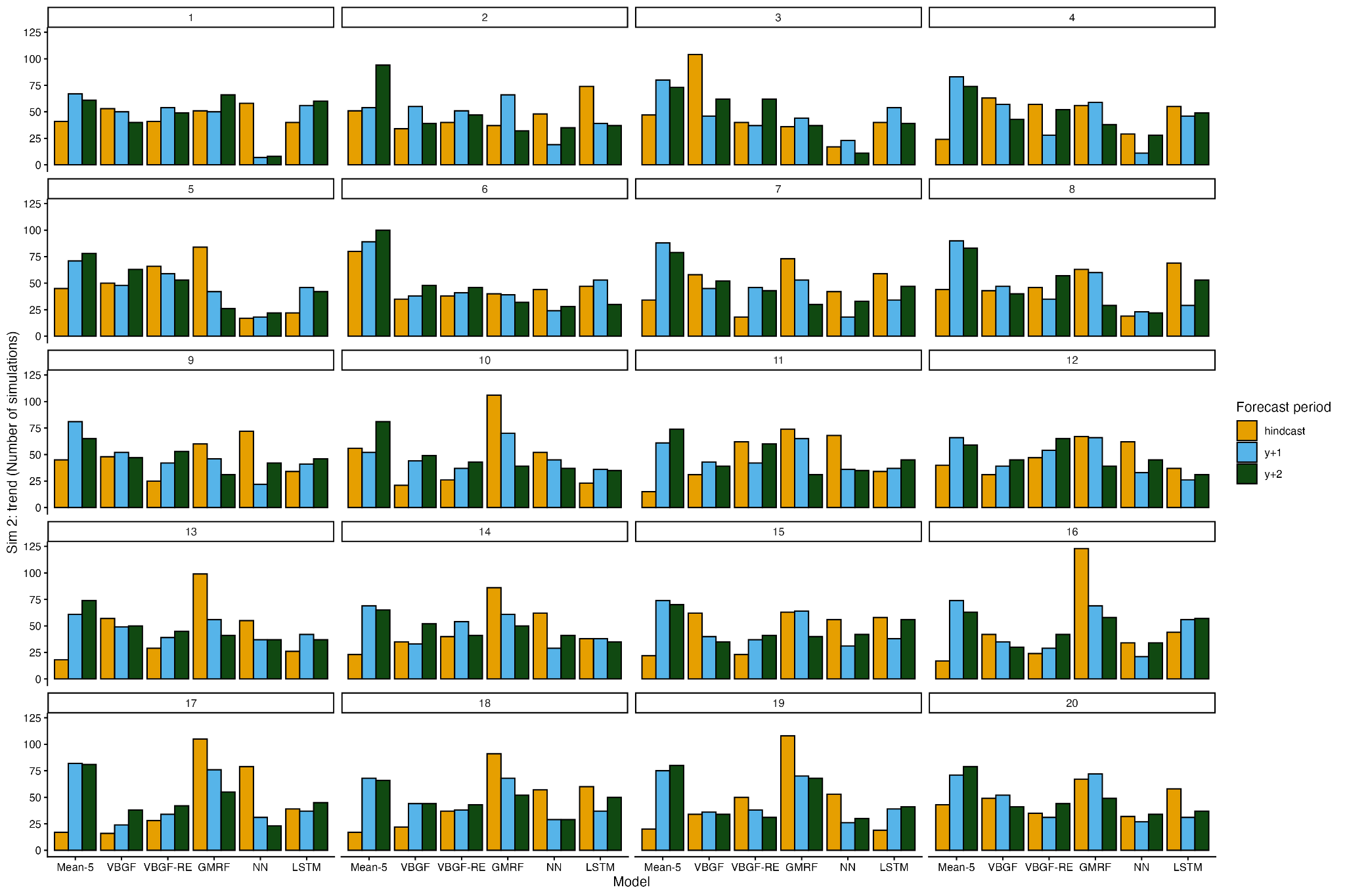

**Figure S2.** Case Study 1: Number of times each process- and neural net based model resulted in the lowest average root mean squared error (RMSE) per age when fit to time-varying simulated data (n = 300). Hindcast represents 10-fold leave year out cross-validation and “y+1” and “y+2” represents forecast skill from sequential peels of historical data and forecasted for two future years.

Supplementary Material S.1. Equations for simulating size-at-age.

Weight-at-age was simulated from a modified von-Bertalanffy growth equation:

$$W_{i}=W_{\infty,y}\left( 1-e^{-K_{y}\left( age-t_{0,y} \right)} \right)e^{\varepsilon_{i}}$$

$$\boldsymbol{\theta}_{\boldsymbol{y}}=\left[ ln\left( W_{\infty,y} \right),ln\left( K_{y} \right),t_{0,y} \right]$$

$$\mu_{y,p}=\bar{\mu}_{p,sim}\left( 1+\beta_{p}*\frac{y-1}{T-1} \right)$$

$$\boldsymbol{\theta}_{\boldsymbol{y}}\boldsymbol{\sim}\mathrm{MVN}\left( \boldsymbol{\mu}_{\boldsymbol{y}}\boldsymbol{,}\boldsymbol{\Sigma} \right)$$

$$\bar{\mu}_{p,sim}\boldsymbol{\sim}\mathrm{MVN}\left( \bar{\mu}_{p}\boldsymbol{,}\boldsymbol{\Sigma}_{\mathbf{p}} \right)$$

$$\boldsymbol{\Sigma=V\bigotimes C}$$

$$C_{i,j}=\rho^{|i-j|}$$

Where $W_{i}$ is the observed weight-at-age, $W_{\infty,y}$ is the year-specific asymptotic weight, $K_{y}$, is a year specific growth coefficient, $t_{0,y}$ is the year-specific theoretical age at weight zero, and $\varepsilon_{i}$ is the lognormally distributed error. Year specific log asymptotic weight ($ln\left( W_{\infty,y} \right)$), log growth coefficient ($ln\left( K_{y} \right)$), and theoretical age at weight zero are derived from a multivariate normal distribution where the annual mean ($\theta_{y,p}$) of each parameter *p* follows parameter specific trend specified by $\beta_{p}$ multiplied against mean von-Bertalanffy growth paremeters ($\bar{\mu}_{p}$) . The standard deviation of the multivariate normal distribution is the Kronecker product of a variance-variance matrix describing the annual variation in the von-Bertalanffy growth paremeters ($\mathbf{V)}$ and an AR1 process **(**$\mathbf{C}$**).**

For all simulations, the maximum age was selected to be 20 and the number of annual samples 300. Mean von-Bertalanffy growth parameters ($\bar{\mu}_{p,sim}$) for each simulation were simulated from the mean ($\bar{\mu}_{p}$) and associated variance-covariance ($\boldsymbol{\Sigma}_{\mathbf{p}}$) derived from FishLife following *Gadus* life history (Thorson et al., 2023). Values for all parameters used in each simulation experiment are described below.

| **Parameter** | **Simulation** | **Value or distribution** |
| --- | --- | --- |
| $ln\left( W_{\infty} \right)$ | 1 and 2 | 8.235583 |
| $ln\left( K \right)$ | 1 and 2 | -1.620628 |
| $t_{0}$ | 1 and 2 | -0.1 |
| $\boldsymbol{\Sigma}_{\mathbf{p}}$ | 1 and 2 | 0.37158203, -0.07989502, 0  -0.07989502, 0.04229736, 0,  0, 0, 0.00250000 |
| $\varepsilon_{i}$ | 1 and 2 | $\varepsilon_{i}\sim\mathrm{LN}\left( \log\left( 0.15 \right), 0.5 \right)$ |
| $\rho$ | 1 | 0 |
| $\beta_{p}$ | 1 | 0, 0, 0 |
| $\mathbf{V}$ | 1 | 0, 0, 0  0, 0, 0,  0, 0, 0 |
| $\rho$ | 2 | $\rho\sim U\left( 0.8, 0.95 \right)$ |
| $\beta_{p}$ | 2 | $\boldsymbol{\beta\sim}\mathrm{MVN}\left( 0,\Sigma_{\beta} \right)$ |
| $\Sigma_{\beta}$ | 2 | 0.01000000, -0.00637288, 0  -0.00637288, 0.01000000, 0  0, 0, 0 |
| $\mathbf{V}$ |  | 0.0412868924, -0.0088772244, 0  -0.0088772244 0.0046997070, 0  0, 0, 0.0002777778 |

Thorson, J.T., Maureaud, A.A., Frelat, R., Mérigot, B., Bigman, J.S., Friedman, S.T., Palomares, M.L.D., Pinsky, M.L., Price, S.A., Wainwright, P., 2023. Identifying direct and indirect associations among traits by merging phylogenetic comparative methods and structural equation models. Methods Ecol. Evol. n/a. <https://doi.org/10.1111/2041-210X.14076>

Supplementary Material S.2 Details on the Reinforcement Learning Case Study

### Management Strategy Evaluation for Eastern Bering Sea Pollock

We first conducted an MSE of the EBS pollock fishery that implemented the currently-used harvest control rule. The MSE was based on an age-structured assessment model constructed in Rceattle that is parameterized similarly to the Eastern Bering Sea pollock assessment model used for tactical management advice ([Adams et al., 2025](https://cdnsciencepub.com/doi/10.1139/cjfas-2024-0225)). The operating model (OM) was conditioned to historical survey and fishery data on age- and length-composition, indices of relative abundance, and catch between 1964 and 2024. The model was projected forward from 2025 to 2050 assuming constant growth, selectivity, and recruitment according to each of the two regimes described above (constant or Ricker). For each future year, data were resampled assuming the same observation error as the terminal year of the conditioning period and passed to an estimation model with an identical structure as the OM, which returned estimates of stock status. For the MSE, the harvest policy used to determine future catches was the North Pacific Fishery Management Council Tier 3 Harvest Control Rule (NPFMC 2017), which is a rectilinear harvest control rule based on spawner-per-recruit-based fishing mortality rates (F-SPR) and sets spawning biomass (SB) targets to SB at F-SPR-40% and closes the pollock fishery when SB falls below 20% of unfished equilibrium SB. This management strategy did not change between the two recruitment scenarios.

### Reinforcement Learning Agent Specification

We then separately trained an RL algorithm on the historical period (1964-2024) of the EBS pollock operating model to discover a optimal harvest policy based on observed bottom trawl survey biomass, and projected the population forward under that policy for 25 years (2025-2050).

For the two recruitment scenarios, the RL agents were given the same training configurations. At each time step, the agents made observations of the environment via the Bottom Trawl Survey and prescribed actions by proposing annual catches. The reward function – the feedback signal returned to the agent at each time step – was defined as the realized, annual harvested biomass. Therefore, the agent was exclusively rewarded for high catches and was not penalized for low stock biomass; earlier attempts at including a composite reward function with these two quantities produced agents that performed sub-optimally. The Proximal Policy Optimization (PPO) algorithm was used for agent training as it has been shown to achieve good performance across a wide range of sequential decision-making problems without requiring extensive tuning. Following Montealegre-Mora et al. (2025), we used a neural network with three levels to represent the nonlinear harvest control rule (i.e., the policy function that prescribes catch based on observed survey biomass)^[1]^. Agents were trained with a budget of 2 million timesteps. Training was performed on 25 stochastic evaluation replicates (random recruitment deviates) on a desktop computer with 24 core CPU with 128 GB of RAM and completed within 24 hours.

1. For brevity, we do not describe the role of the value function in RL. See (Schulman et al., 2017).
